# SOL1 and SOL2 Regulate Fate Transition and Cell Divisions in the Arabidopsis Stomatal Lineage

**DOI:** 10.1101/394940

**Authors:** Abigail R. Simmons, Kelli A. Davies, Wanpeng Wang, Zhongchi Liu, Dominique C. Bergmann

## Abstract

In the stomatal lineage, cells make fate transitions from asymmetrically dividing and self-renewing meristemoids, to commitment to the guard mother cell identity, and finally though a single division to create mature, post-mitotic stomatal guard cells. Flexibility in the stomatal lineage allows plants to alter leaf size and stomatal density in response to environmental conditions; however, transitions must be clean and unidirectional in order to produce functional and correctly patterned stomata. Among direct transcriptional targets of the stomatal initiating factor, SPEECHLESS, we found a pair of genes, *SOL1* and *SOL2*, required for effective transitions in the lineage. Here we show that these two genes, which are homologues of the LIN54 DNA-binding components of the mammalian DREAM complex, are expressed in a cell cycle dependent manner and regulate cell fate and division properties in the self-renewing early lineage. In the terminal division of the stomatal lineage, however, these two proteins appear to act in opposition to their closest paralogue, *TSO1*, revealing complexity in the gene family may enable customization of cell divisions in coordination with development.

## Introduction

The development of organized tissues containing multiple cell types requires a careful balance of proliferation and differentiation processes. One such balancing act is found in the leaves of *Arabidopsis*, where divisions in the stomatal lineage generate the majority of epidermal cells (Geisler et al., 2000). The stomatal lineage is characterized by an early proliferative meristemoid phase in which cells divide asymmetrically in a self-renewing fashion, followed by a transition and commitment to one of two alternative fates: pavement cell or guard mother cell (GMC). If a cell becomes a GMC, it will divide symmetrically to form the two guard cells of the stomatal complex, a valve-like structure that facilitates plant/atmosphere gas exchange (Fig. 1A).

**Figure 1:**
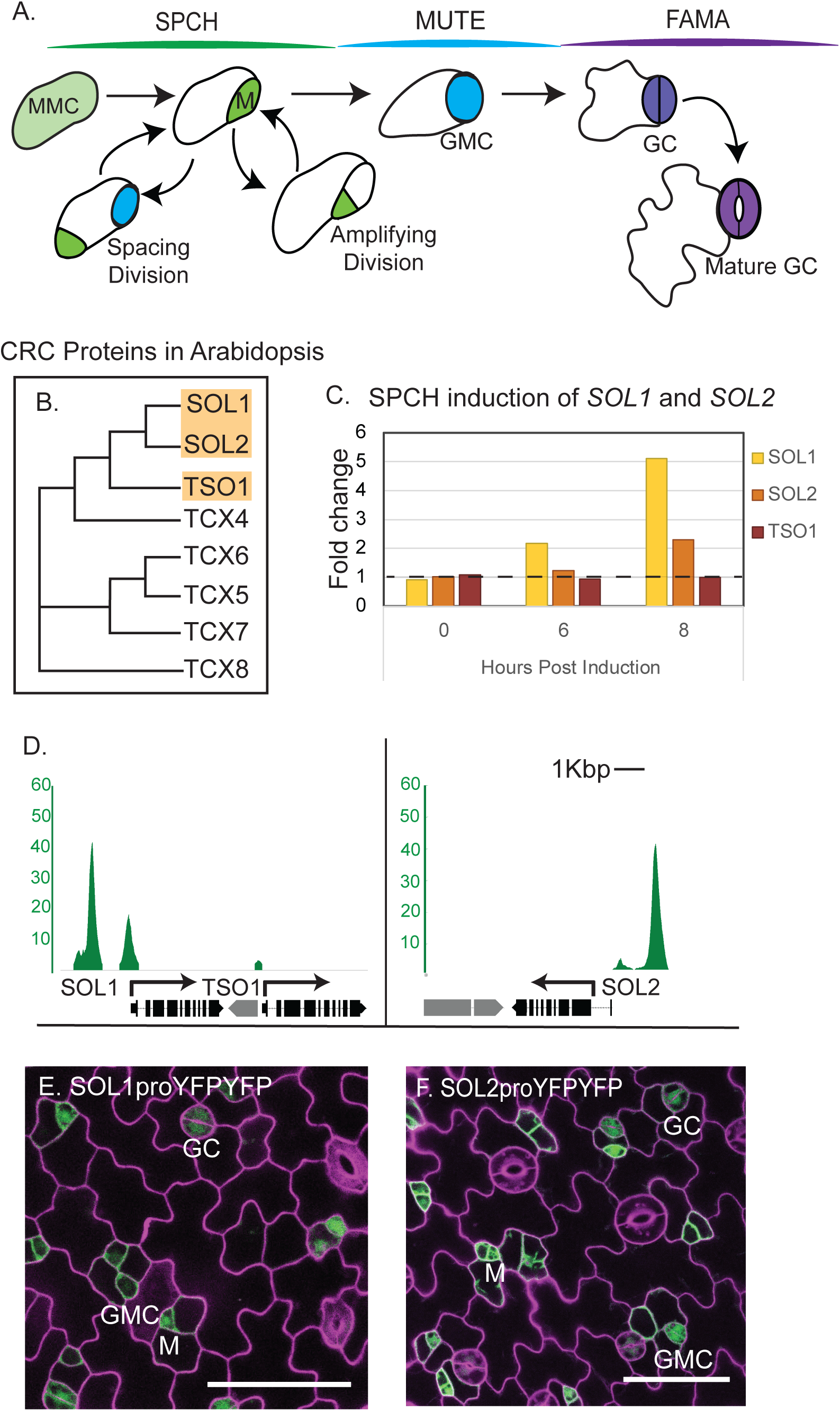
SPCH targets *SOL1* and *SOL2* are expressed in the stomatal lineage. **(A)** Schematic of stomatal development; each stage color-coordinated with the bHLH transcription factor that regulates it: SPCH (SPEECHLESS) in meristemoid (M) phase, MUTE in guard mother cell (GMC) phase, and FAMA in the guard cell (GC) differentiation phase. **(B)** Phylogenetic tree of CHC proteins in Arabidopsis, with subjects of this paper shaded, produced with Clustal Omega. **(C)** Evidence that *SOL1* and *SOL2* transcripts increase in response to estradiol induction of SPCH; fold change over estradiol induced wildtype control (Lau et al. 2014). **(D)** SPCH ChIP-seq reveals promoters of SOL1 and SOL2 are bound by SPCH; y-axis represents enrichment value (CSAR), the output score from MACS2, in arbitrary units from (Lau et al. 2014). **(E-F)** Confocal images of *SOL1* and *SOL2* transcriptional reporters (green) in 3 dpg abaxial cotyledon, indicating they are expressed in meristemoids (M), guard mother cells (GMCs) and young guard cells (GC). Cell outlines (purple) visualized by staining with propidium iodide. 50 μm scale bars.

**Figure 2:**
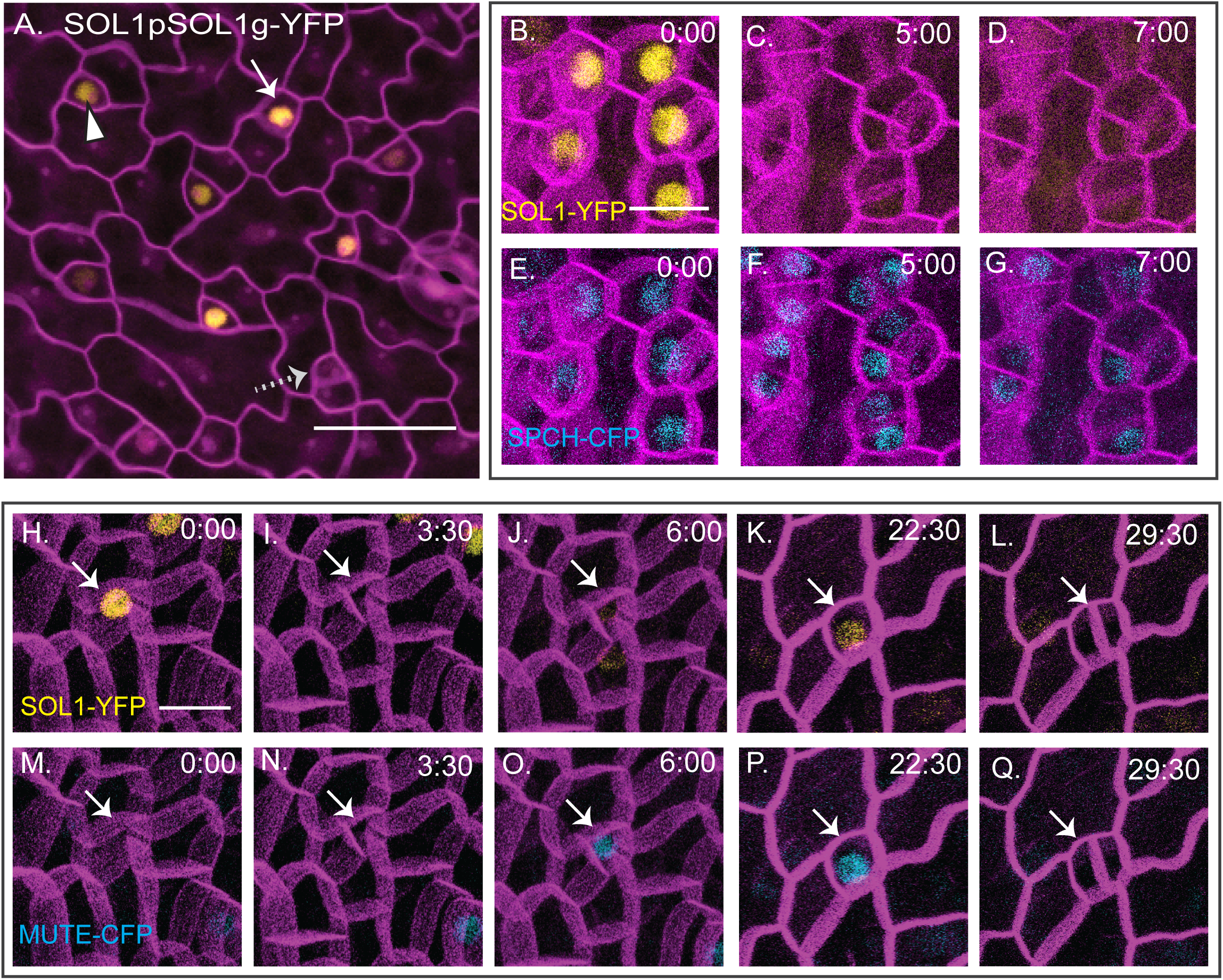
SOL1 is co-expressed with SPCH and MUTE prior to asymmetric and symmetric divisions. **(A)** A functional SOL1-YFP reporter is expressed in some (white arrow), but not all (dashed arrow) meristemoids and GMCs (arrowhead) in 3 dpg abaxial cotyledons (full genotype: SOL1p:SOL1-YFP; *sol1 sol2*); cell outlines visualized with propidium iodide (purple); scale bar 50 μm. **(B-Q)** Time-lapse confocal images, cell outlines (purple) visualized with ML1pro:RCI2A-mCherry in wildtype background, time in hour:minutes noted in top right of each image, scale bars 10 μm. **(B-G)** Time-lapse of SOL1pro:SOL1-YFP (yellow, B-D) and SPCHpro:SPCH-CFP (blue, E-G). **(H-Q)** Time-lapse of SOL1pro:SOL1-YFP (yellow, H-L) and MUTEproMUTE-CFP (blue, M-Q). Arrows follow a single cell through an asymmetric division (I and N), conversion to round GMC (K and P) and a symmetric division generating paired guard cells (L and Q).

Transcriptional regulation of division and differentiation in the stomatal lineage involves a set of closely related and sequentially expressed basic helix loop helix (bHLH) transcription factors, SPEECHLESS (SPCH), MUTE and FAMA (Fig. 1A) and their more distantly related bHLH heterodimer partners ICE1/SCREAM and SCRM2. These transcription factors regulate both cell fate and cell division. For example, in the ultimate product of the stomatal lineage, guard cells, RETINOBLASTOMA RELATED (RBR) is needed to halt divisions (Borghi et al., 2010) and also forms a complex with FAMA to maintain mitotic quiescence and keep guard cells in a terminally differentiated state (Lee et al., 2014; Matos et al., 2014). FAMA also directly represses cell-type specific CYCLIN(CYC) D7;1 to prevent over-division of guard cells (Weimer et al., 2018). One stage earlier, MUTE is required to repress the previous meristemoid fate and simultaneously drive cells to adopt GMC fate (Pillitteri et al., 2007). MUTE does so in part by directly regulating CYCD5;1 and other cell cycle factors to ensure the GMC divides symmetrically to form the guard cells (Han et al., 2018).

The earliest phases of the stomatal lineage are complicated because there are three types of asymmetric divisions--entry, amplifying and spacing--that occur an indeterminate number of times. Previous studies have sought to understand how SPCH controls entry into the stomatal lineage and how SPCH drives these recurrent and varied asymmetric divisions. From these studies, positive and negative feedback motifs emerged, with SPCH inducing its transcriptional partners ICE1 and SCRM2 to locally elevate its activity, while also initiating a longer range negative feedback through secreted signaling peptides to ensure its eventual downregulation (Horst et al., 2015; Lau et al., 2014). Targets that connect SPCH to core cell cycle behaviors and that allow meristemoids to exit the self-renewing stage and progress to GMCs, however, remained elusive.

Here we characterize the expression pattern and function of *SOL1* and *SOL2*, two genes encoding proteins containing cysteine rich-repeat (CXC) domains separated by a conserved hinge (<u>C</u>XC-<u>H</u>inge-<u>C</u>XC, CHC), in the stomatal lineage. Their expression patterns are not identical, but both genes are enriched in the stomatal precursors, and protein reporters accumulate in nuclei in a distinct pattern coincident with cell-cycle progression. We show the SOL1 and SOL2, although initially identified as SPCH target genes, are required for efficient fate transitions through multiple stomatal lineage stages and in their absence, cell fates are incorrectly specified. Finally, we consider a potentially antagonistic relationship between these two genes and their next closest paralogue, *TSO1*, in the final guard-cell generating division of the stomatal lineage.

## Results

### SOL1 and SOL2 are stomatal-lineage expressed targets of SPCH

Among the hundreds of genes both bound and upregulated by SPCH, we were particularly drawn to two genes encoding CHC proteins. Animal CHC proteins LIN54 (*C. elegans, H. sapiens*) and MYB interacting protein (MIP) 120 (*D. melanogaster*) bind DNA in a sequence specific manner and are components of DREAM (DP, RBR, E2F and Myb-MuvB (Multi-vulval Class B)) complexes. Animal DREAM complexes are implicated in cell-cycle and transcriptional regulation, chromatin remodeling and cell differentiation (Sadasivam and DeCaprio, 2013). Arabidopsis encodes eight CHC-domain proteins ((Andersen et al., 2007); Fig. 1B); of this family, only *TSO1* has been functionally characterized, and *TSO1* is important for properly regulating divisions in the floral meristem (Song et al., 2000). SPCH directly targets At3g22760 and At4g14770 (Fig. 1C-D), a closely related pair located in the same branch of the CHC family as *TSO1.* In the literature, At3g22760 and At4g14770 have been given the names *SOL1/TCX3* (TCX = TSO1-like CXC, SOL = TSO1-like) and *SOL2/TCX2*, respectively (Andersen et al., 2007; Liu et al., 1997; Sijacic et al., 2011). We will refer to these genes as *SOL1* and *SOL2*. *SOL1* and *TSO1* are tandemly arranged in the genome, but *TSO1* does not appear to be a SPCH target (Fig. 1C-D).

To determine the expression pattern of *SOL1* and *SOL2*, we generated transcriptional reporters containing 2457bp and 2513 kb of 5’ sequence, respectively, driving expression of yellow fluorescent protein (YFP). Both *SOL1* and *SOL2* reporters were expressed in young leaves and were most strongly expressed in young stomatal lineage cells, consistent with *SOL1* and *SOL2* being targets of SPCH (Fig. 1E-F). To gain insight into SOL protein behaviors, we generated translational reporters; downstream of the promoters, we added the genomic fragments of SOL1 and SOL2 encompassing exons and introns from the predicted translational start codon to before the stop codon (2757bp genomic and 3301bp respectively) with a 3’ sequence encoding YFP. Both translational reporters were restricted to nuclei (Fig. 2 and Fig. 3) and both appeared to be functional as they rescued the *sol1 sol2* mutant phenotypes in the stomatal lineage (described below and in Fig. 4).

**Figure 3:**
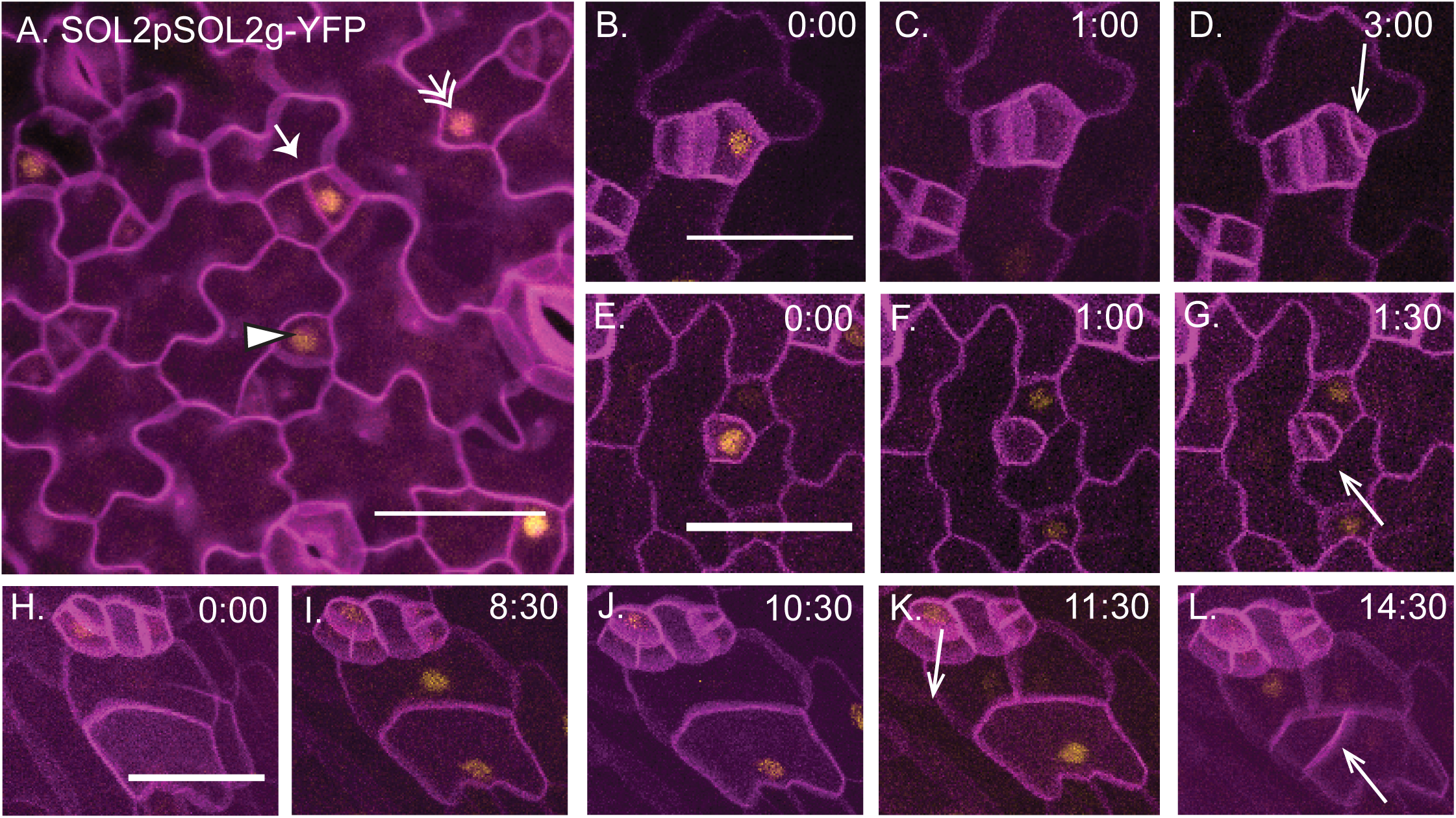
SOL2 is expressed in meristemoids, GMCs and pavement cells in a cell cycle dependent manner. **(A)** A functional SOL2-YFP reporter is expressed in meristemoids (arrow), GMCs (arrowhead) and SLGCs (double arrow) in 3 dpg abaxial cotyledon (full genotype: SOL2p:SOL2-YFP; *sol1 sol2*). Cell outlines stained with propidium iodide (purple); scale bar 50 μm. **(B-L)** Time-lapse images of SOL1pro:SOL1-YFP (yellow) with cell outlines marked by ML1pro:RCI12A-mCherry (purple) time in hour:minutes noted in top right of each image. Arrows indicate new cell divisions. **(B-D)** meristemoid divides asymmetrically. **(E-G)** GMC divides symmetrically. **(H-L)** Pavement cells divide. In each division SOL2 expression disappears 1-2 hours before cell division. Scale bars 50 μm.

**Figure 4:**
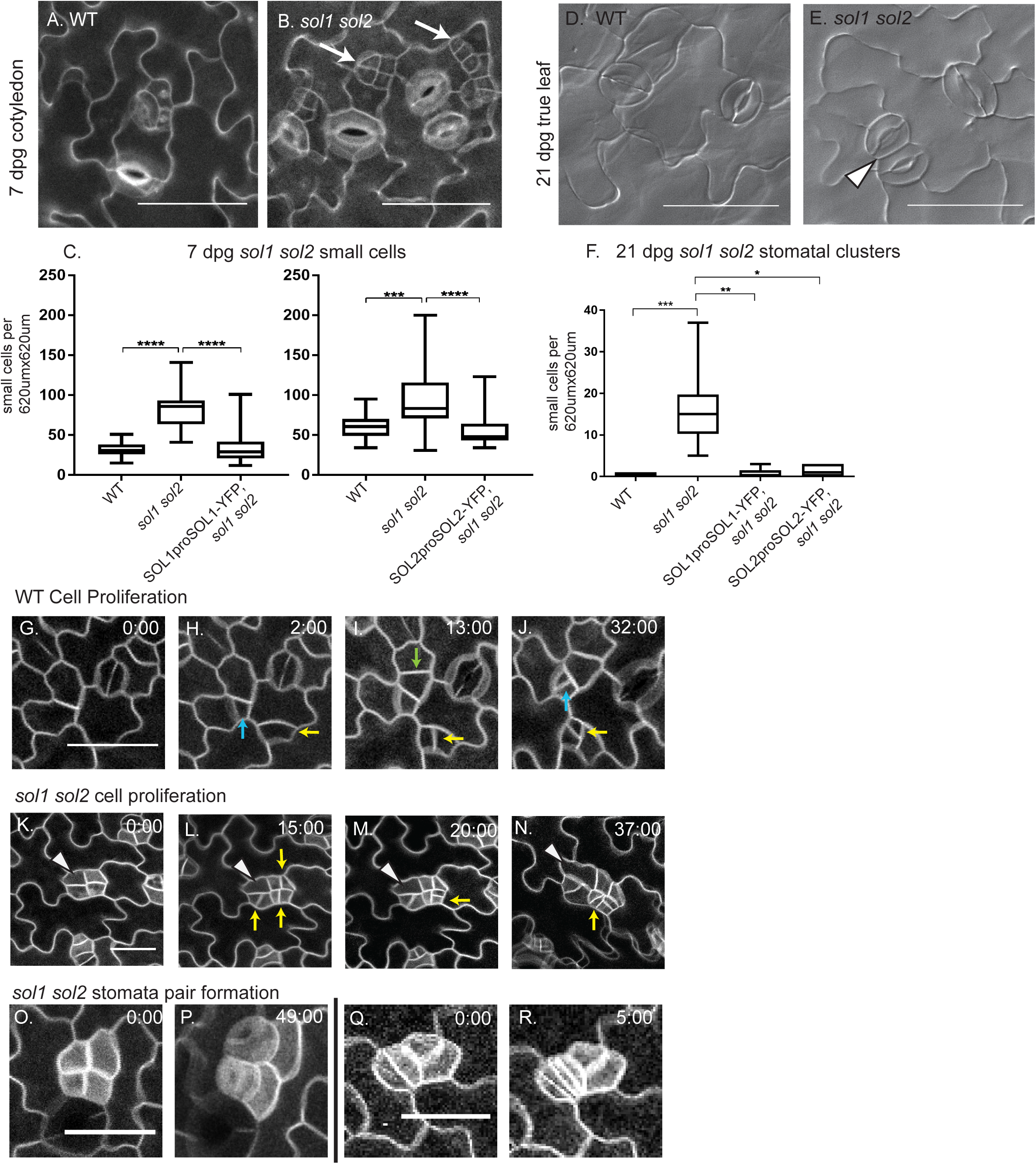
*SOL1* and *SOL2* are redundantly required for control of early and late stomatal cell division behaviors. **(A)** Confocal images of 7 dpg WT abaxial cotyledon containing few small cells (indicated by white arrows) in comparison to **(B)** *sol1 sol2* double mutants. **(C)** Quantification of small cell phenotype, n = 16-22. **(D-E)** DIC images of 21 dpg adaxial true leaf in WT (D) and *sol1 sol2* (E); stomatal pairs indicated with arrowhead. **(F)** Quantification of pairs and higher order stomatal clusters, n = 5-10. (A,B,D,E) 50 μm scalebars. **(G-R)** Time-lapse confocal imaging; cell outlines visualized with ML1pro:RCI2A-mCherry. **(G-J)** Cell proliferation in WT, divisions marked with yellow, blue and green arrows. **(K-N)** Small cell divisions in *sol1 sol2*, cell divisions marked with yellow arrow. One small cell (white arrowhead) begins to lobe. **(O-P)** Two neighboring small cells both divide into stomata. **(Q-R)** Two guard cells each divide symmetrically again. (O, Q) 30 μm scale bars. Significance indicated: * p<0.05, **p<0.01, ***p<0.001, ****p<0.0001. Dunn’s multiple comparison test.

SOL1-YFP was expressed in the meristemoids and GMCs (Fig. 2A). Compared to the corresponding transcriptional reporter, SOL1-YFP showed a somewhat patchy expression pattern. Although it was expressed in nuclei of both GMCs and meristemoids, the brightness varied among populations of these cells (Fig. 2A) and some young stomatal lineage cells did not express it at all (Fig. 2A, dotted arrow). Given the role of SOL1 homologues in the cell cycle, we hypothesized that variation in expression was due to cell-cycle regulated protein abundance. To test this, we performed time-lapse confocal microscopy on SOL1-YFP expressing plants. We included either SPCH-CFP (meristemoid marker) or MUTE-CFP (GMC marker) and a plasma-membrane marker (RCI2A-mCherry) in the background to allow us to precisely identify the cells in which SOL1 was expressed.

SOL1 was co-expressed with SPCH prior to asymmetric divisions of meristemoids (Fig. 2B,E), however the SOL1-YFP signal disappears at the division, while SPCH-CFP persists initially in both daughter cells (Fig. 2C,F), before being retained in only the smaller of the two daughter cells (Fig. 2D,G). Time-lapse imaging of SOL1-YFP and MUTE-CFP shows a similar pattern. Because SOL1-YFP is expressed in meristemoids, it initially precedes MUTE expression (Fig. 2H,M) but disappears before the cell divides (Fig. 2 I,N). In cells transitioning to GMC fate, MUTE-CFP precedes SOL1-YFP expression (Fig.2 J,O), but eventually the two markers are co-expressed (Fig.2 K,P) and both markers are gone prior to the symmetric GMC division (Fig.2 L,Q). Altogether, SOL1 is expressed in nuclei of cells at the early meristemoid stage, the late meristemoid stage and the GMC stage, but it disappears prior to cell divisions, suggesting that the protein is actively degraded in a cell-cycle dependent manner (see also Fig. S1A-E).

SOL2-YFP resembles SOL1-YFP in its co-expression with SPCH-CFP in nuclei of meristemoids and MUTE-CFP in nuclei of GMCs (Fig. S1F,G). SOL2, however, was often also expressed in the sister cells of meristemoids (stomatal lineage ground cells, SLGCs) and in pavement cells (Fig. 3A, double arrows). This expression pattern could emerge from a more broadly expressed promoter, or because SOL2 is under different cell-cycle regulation than SOL1, and simply persists into these cell types after meristemoid division. Time-lapse imaging of SOL2-YFP revealed that expression disappears prior to asymmetric meristemoid divisions (Fig. 3B-D) and symmetric GMC divisions (Fig. 3E-G), just like SOL1. The expanded domain of SOL2 instead appears to be due to expression beginning in pavement cells prior to their division (Fig. 3H-L), just as it does in meristemoids and GMCs. To further narrow down when in the cell cycle SOL2 was expressed, we time-lapse imaged plants co-expressing SOL2-YFP and the S-phase marker HTR2pro:CDT1a(C3)-RFP (Yin et al., 2014). SOL2-YFP was visible on average 3 hours before the CDT1a-RFP (Fig. S1H-L, quantified in M). SOL2 then disappeared 1-2 hours before appearance of the new cell plate; timing that is consistent with degradation during the G2-M transition (Fig. S1N). Taken together, these data suggest that SOL1 and SOL2 could function in the late G1, S, and G2 cell cycle phases in meristemoids and GMCs.

### SOL1 and SOL2 are redundantly required for stomatal lineage progression and correct stomatal patterning

To explore the function of these proteins in the stomatal lineage, we identified T-DNA insertion alleles for each and tested their impact on *SOL1* or *SOL2* expression (Fig. S2A). Two alleles for each gene dramatically reduced expression as assayed by qRT-PCR, though none completely abolished it (Fig. S2B). Double mutants were generated by crossing and genotyping for the relevant mutation by PCR (details in methods). A typical phenotype for disruptions in stomatal lineage cell fate, signaling or polarity is the presence of stomata in pairs or clusters in mature cotyledons, so we counted stomatal pairs on 21 days post germination (dpg) adaxial cotyledons for each single mutant and two double mutant combinations. No *SOL1* or *SOL2* single mutants had a statistically significant pairing phenotype, but both double mutant combinations did (Fig. S2C). The strongest pairing phenotype and lowest expression of *SOL1* and *SOL2* genes was found in the *sol1-4 sol2-2* double mutant, and so we focused on this double mutant for more detailed phenotypic analysis; unless otherwise mentioned, *sol1 sol2* will refer to this specific allelic combination.

To capture the complexity of divisions and fates in the stomatal lineage, we characterized the *sol1 sol2* phenotype at 7 dpg, when SPCH-associated amplifying divisions are occurring, and a late stage (21 dpg) when the (wildtype) epidermis has finished development and contains only mature guard and pavement cells. At 7 dpg in abaxial cotyledons, the most distinctive *sol1 sol2* phenotype was the increased number of small cells (here defined as cells less than 200 square micron in area), often found in clusters (Fig. 4B, white arrows). Wildtype seedlings have some of these small cells (Fig. 4A), however, the number is significantly increased in *sol1 sol2* double mutants (Fig. 4B-C) and this small cell phenotype can be rescued by expression of SOL1 or SOL2 reporters (Fig. 4C).

We next examined the end stage phenotype of the first pair of true leaves at 21 days post germination (dpg). In wildtype seedlings, the adaxial true leaf epidermis consists mostly of guard cells and pavement cells (Fig. 4D). In *sol1 sol2* double mutants at this stage, the most prominent phenotype was pairs of stomata (Fig. 4E, white arrowhead). Resupplying SOL activity via translational reporter also rescued this late stage phenotype (Fig. 4F). We chose to score the adaxial true leaf as representative of an end stage phenotype, because cells in the abaxial true leaf in *sol1 sol2* mutants were still dividing at 21 dpg, a phenotype in itself. Both abaxial and adaxial true leaves, however, contained stomatal pairs at this late stage.

We used time-lapse imaging to pinpoint the origin of the early and late stomatal lineage phenotypes and the connection between them. A key question is whether the accumulation of small cells comes from aberrant divisions (e.g. divisions of non-stomatal lineage cells, or inappropriately symmetric divisions) or whether divisions are qualitatively normal, but more frequent. *sol1 sol2* cotyledons marked with plasma membrane marker ML1pro:RCI2A-mCherry were tracked for 60 hrs (images captured every 60 min, starting age 3 dpg when the stomatal lineage is initiating), and compared to a time matched series from a wildtype cotyledon. Stomatal lineage progression is asynchronous, and we followed cells from regions displaying a diversity of mature and precursor cell types.

In wildtype, we observed frequent asymmetric divisions of meristemoids (Fig. 4H, yellow and blue arrows). The asymmetrically dividing meristemoid cells appeared, in the plane of the epidermis, as slightly lobed squares, and typically divided 1-2 more times in a spiral pattern previously described as “amplifying divisions” (Geisler et al., 2000; Robinson et al., 2011) (Fig. 4I-J, yellow arrows). Visually symmetric divisions were also observed in larger cells (Fig. 4I, green arrow).

In *sol1 sol2* mutants, we also observed repeated divisions of slightly lobed square cells (Fig. 4K-N, Fig. S3A-D) and while it was clear that the mutant seedlings had more small cells than wildtype, our data did not suggest that the small cells resulted from qualitatively aberrant divisions. For example, in Fig. 4K-L, three of the four small cells undergo an asymmetric division, each of which appears normal in terms of size and orientation. Some of the small cells generated in this manner continued in the lineage, ultimately dividing symmetrically and forming stomata (Fig. 4N, yellow arrow), but others remained small during the time course. One of the cells, (Fig. 4K-N, white arrowhead) did not divide in the course of the video and instead began to form lobes. In other cases, groups of four small cells were observed to arise from additional divisions of a meristemoid/SLGC pair (e.g. Fig. S3E-G).

Since the early asymmetric divisions appeared qualitatively normal, we considered alternative explanations for the appearance of excess small cells: cells might divide faster or post-division expansion could be slowed. To evaluate these possibilities, we needed to be able to monitor a cell from its initial “birth” until its next division, which was challenging due to the typical (>16hr) length of plant cell cycles, but from the time-lapse movies we were able to quantify 24 such divisions in WT and 22 divisions in *sol1 sol2*. We calculated cell cycle length as the time (in hours) between one cell division and the next, and areal expansion as the traced 2D area of a cell immediately after its first division compared to immediately before its second division. We found that the cell cycle in *sol1 sol2* double mutants was significantly *slower* than in wildtype (4.5 hours median difference, Fig. S3H). The percent areal growth per hour however, was also significantly less (Fig. S3I and methods). Overall leaf size in *sol1 sol2* was not significantly different from wildtype at 14 dpg (Fig. S3J), consistent with the smaller cell size balancing out the effect of greater cell numbers observed in the mutants. Failure to expand post division is a hallmark of cell identity defects in SLGCs and can be seen when SPCH or ICE1 are not correctly degraded in SLGCs (Kanaoka et al., 2008; Lampard et al., 2008). When SPCH or ICE1 is stabilized, SLGCs maintain the division capacity of their SPCH-expressing predecessors, leading to the accumulation of excess small cells.

### SOL1 and SOL2 activity appears to be required at multiple transitions

The late-stage phenotype of stomatal pairs could arise from inappropriate divisions of GCs, or from earlier defects such as cell identity errors in SLGCs that enable both these cells and their sister cells to act as guard cell precursors. When we extracted examples of stomatal pair formation from the time-lapse images, we observed two origins for stomatal pairs. In some cases, two small cells in a group of four differentiated into GMCs and then divided to form stomata in contact (Fig. 4O-P); showing that the early stage phenotype can develop directly into the late stage phenotype. However, we also observed two young guard cells both divide a second time to produce four guard cells (Fig. 4Q-R) suggesting *SOL* activity at the MUTE stage or later was required. These two defects suggest multiple roles for SOLs in stomatal transitions and are consistent with the expression of SOLs just prior to the meristemoid division and the GMC division.

### MUTE expression is disconnected from cell fate in *sol1 sol2* double mutants

Division behaviors suggested cell identity defects in the stomatal lineage, but to more accurately characterize these defects, we examined SPCH, MUTE and FAMA translational reporters in *sol1 sol2* mutants. To capture the very earliest stages of the lineage, we imaged cotyledons at 3 dpg as well as at 7 dpg. SPCH is expressed in small cells in *sol1 sol2* and wildtype at 3 dpg (Fig. 5A and Fig S4A), though there are more of these small cells in the mutant. At 7 dpg, small cells that have begun to lobe lose SPCH (Fig. 5B), suggesting that the small cells are likely meristemoids and that SPCH is not obviously mis-regulated in the absence of *SOL1* and *SOL2*. A similar comparison of MUTE expression at these two timepoints did reveal a deviation from WT in that the number of cells expressing MUTE did not decrease over time (Fig. 5C-D). Because elevated MUTE can lead to stomatal hyperproduction (Pillitteri et al., 2007), we also imaged a transcriptional reporter (MUTEpro:CFPnls) in addition to MUTEpro:MUTE-CFP to confirm that MUTE persistence was not due to the effect of an additional copy of MUTE (Fig. S4G,H). FAMA is mostly expressed in recently divided guard cells at 3 and 7 dpg, but is occasionally observed in rounded small cells that are likely to divide symmetrically (Fig. 5E,F), suggesting that most small cells in *sol1 sol2* have not entered the later (FAMA) stage of the lineage (wildtype comparisons for all markers in Fig. S4).

**Figure 5:**
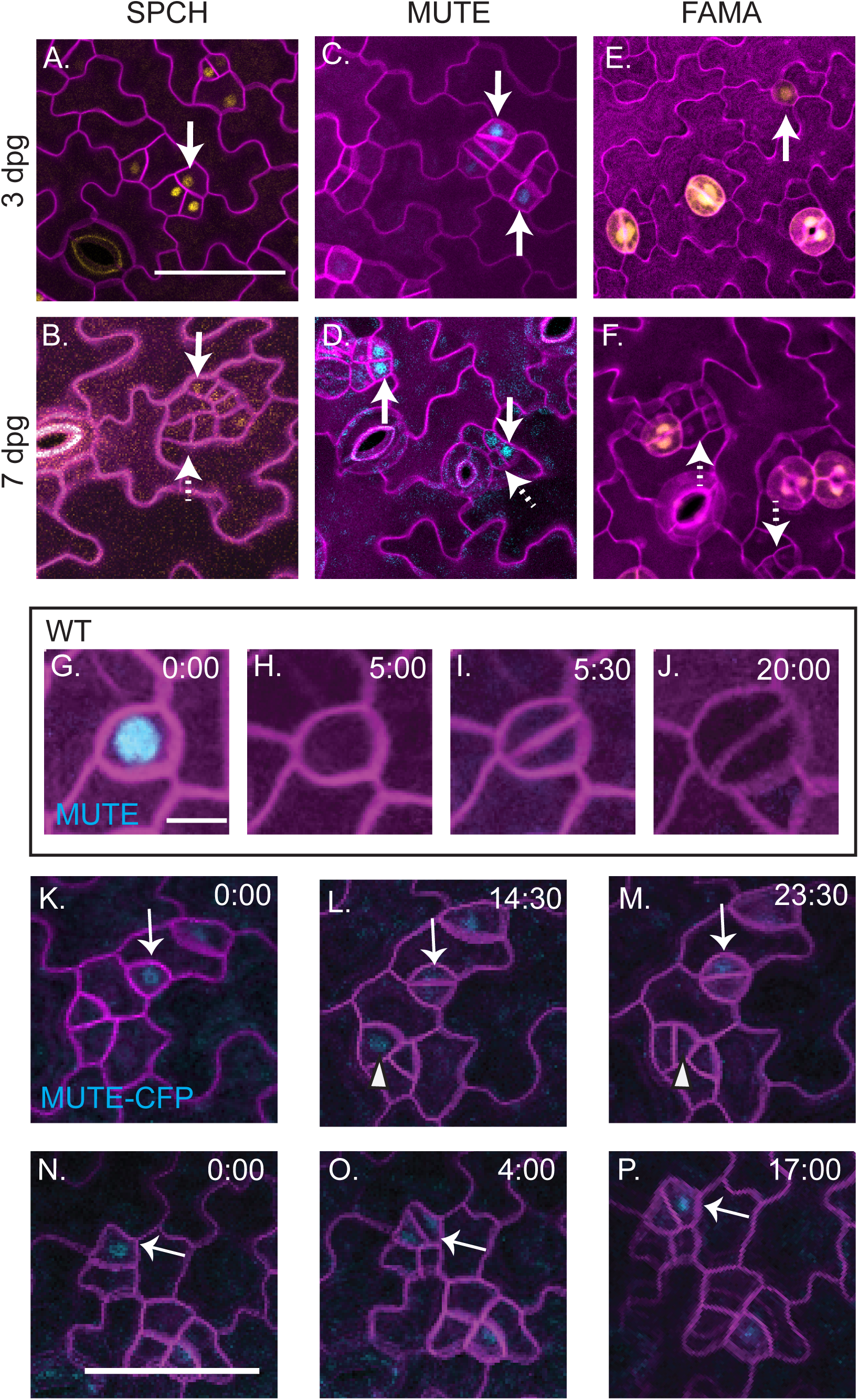
Markers of cell fate are inappropriately expressed in *sol1 sol2* mutants. **(A-F)** Confocal images of abaxial cotyledons from *sol1 sol2* mutants, at indicated days post germination, with cell fate reporters SPCHpro:SPCH-YFP (A-B), MUTEpro:MUTE-CFP (C-D) and FAMApro:YFPnls (E-F). Cell outlines (purple) visualized with propidium iodide. All images same scale, scale bar 50 μm. **(G-J)** Selections from time-lapse of ML1pro:RCI2A-mCherry and MUTEpro:MUTE-CFP marker in WT, 10 μm scale bar where MUTE expressing GMC divides symmetrically. (K-M, N-P) selections from time-lapse of *sol1 sol2* mutant expressing ML1pro:RCI2A-mCherry and MUTEpro:MUTE-CFP markers, all images same scale, 30 μm scale bar in N. **(K-M)** Two MUTE expressing cells (indicated by solid white arrow and arrowhead) divide. **(N-P)** MUTE expressing cell (indicated by solid white arrow) divides asymmetrically.

The appearance of MUTE expressing cells at both 3 dpg and 7 dpg timepoints made us curious about whether the MUTE-positive small cells at 3 dpg progress in the lineage to form guard cells, or if they are stuck at an earlier stage. To determine the fate of MUTE expressing small cells, we performed time-lapse imaging on a MUTE-CFP reporter in *sol1 sol2* seedlings (3 dpg abaxial cotyledon). In wildtype plants, MUTE expression begins after the final asymmetric division (Fig. 5G) and it disappears prior to the symmetric division (Fig. 5H, I), thus MUTE expressing cells do not normally divide in wildtype plants. When we performed time-lapse imaging on *sol1 sol2* lines, however, we found that small cells expressing MUTE-CFP often divide. Sometimes these divisions are visually symmetric, like GMC divisions; however, MUTE expression is still detected long after the division (Fig. 5K-M, white arrow). Other divisions resemble asymmetric meristemoid divisions (Fig. 5L-M white arrowhead, N-P white arrow). Thus, in the absence of *SOL1* and *SOL2*, MUTE expression is no longer sufficient to reliably predict GMC fate.

### *SOL1* and *SOL2* may oppose activity of paralogue *TSO1* in the stomatal lineage

SOL1 and SOL2 are closely related to the CHC-domain protein best characterized in plants, TSO1(Andersen et al., 2007; Sijacic et al., 2011). We did not originally focus on *TSO1* because it is neither bound nor induced by SPCH (Fig.1 B,C and Lau et al, 2014), but a recent publication included a TSO1 translational reporter (Wang et al., 2018) and we found this reporter to be expressed in a pattern similar to SOL2-YFP. Specifically, TSO1-GFP was expressed throughout the epidermis, in meristemoids (Fig. 6A, arrow), GMCs (Fig. 6A, arrowheads) and pavement cells (Fig. 6A, double arrow), but not guard cells. This led us to speculate that *TSO1* could be partially redundant with *SOL1* and *SOL2*.

**Figure 6:**
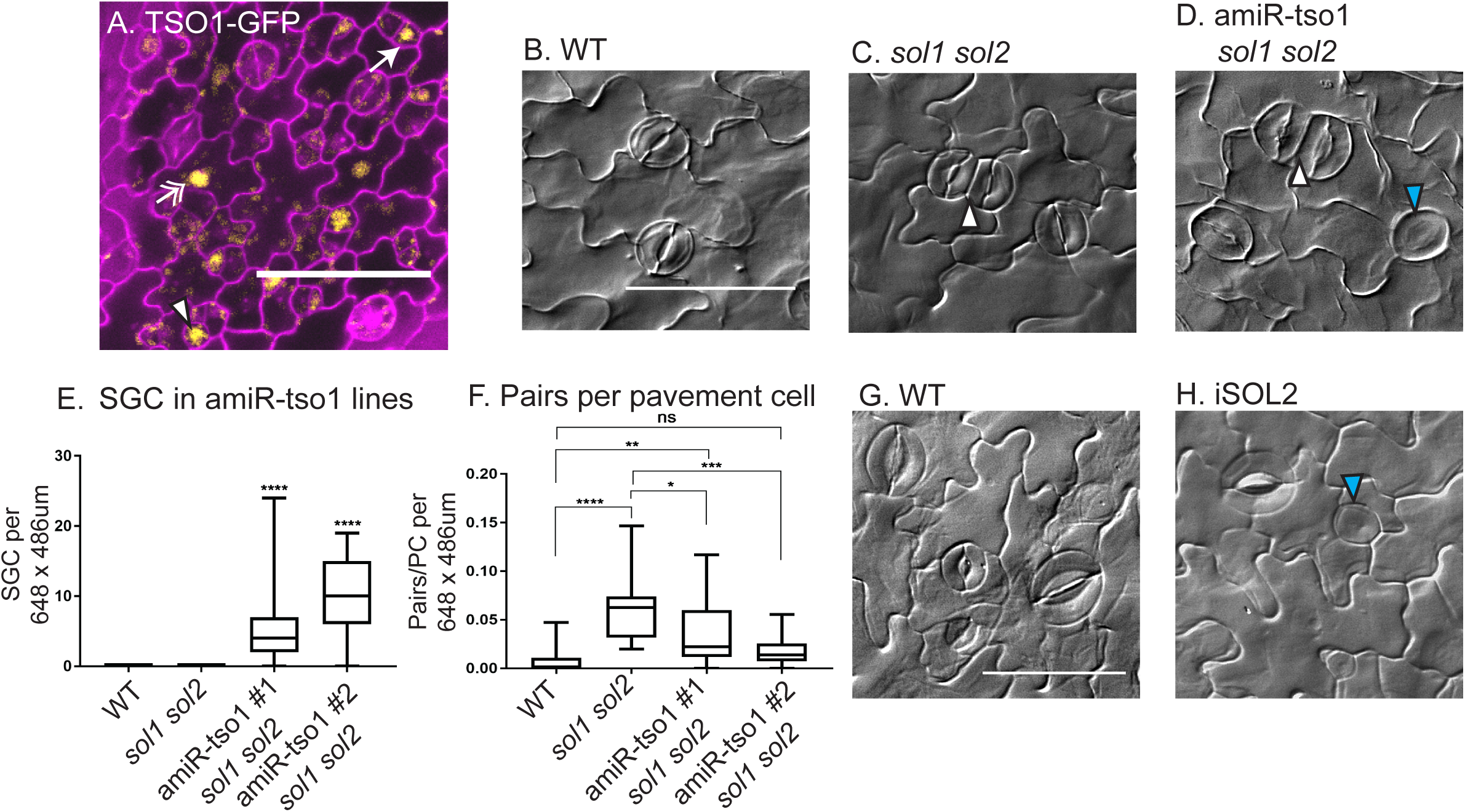
Depletion of *TSO1* in *sol1 sol2* background or overexpression of SOL2 result in similar guard cell division defect. **(A)** Confocal image of TSO1pro:TSO1-GFP reporter expressed throughout epidermis, in meristemoids (arrow), GMCs (arrowhead) and pavement cells (double arrow). **(B-D)** DIC images of 21 dpg adaxial true leaves (B) WT, (C) Stomatal clustering (white arrowhead) in *sol1 sol2*, (D) stomatal pairs (arrowhead) and single guard cells (SGCs, blue arrowhead) in *amiR-tso1 sol1 sol2*. **(E)** Quantification of number of SGCs per field of view. (F) Quantified pairs of stomata per pavement cell in field of view. 50 μm scale bars, n = 19-31. (G-H) DIC images showing production of SGCs upon SOL2-YFP overexpression. Significance indicated: * p<0.05, ** p<0.01, *** p<0.001, **** p<0.0001, Dunn’s multiple comparison test.

The *TSO1* gene is adjacent to *SOL1* (Fig. 1D), which made generating a triple mutant by crossing infeasible, so we reduced expression levels of *TSO1* in the stomatal lineage by expressing an artificial miRNA against it with the *TOO MANY MOUTHS (TMM)* promoter (Nadeau and Sack, 2002). In the *sol1 sol2* background, multiple independent TMMpro:amiRNA-tso1 lines led to an unexpected new phenotype in which guard cells failed to divide, and instead formed large round- or kidney-shaped cells. We termed this phenotype single guard cell, or SGC (Fig. 6D, blue arrowhead), to be consistent with previous literature describing this phenotype (Boudolf et al., 2004; Xie et al., 2010). The SGC phenotype was not described in previous reports on *TSO1* (Andersen et al., 2007; Liu et al., 1997), and our own analysis of segregating populations from two previously described alleles (*tso1* homozygotes are sterile) *tso1-1/sup-5* and SALK_074231C, *tso1-6/+* failed to identify the SGC phenotype (no instances in 18 seedlings from *tso1-1/sup-5* plants and 24 seedlings from *tso1-6*/+). We therefore concluded that in the *sol1 sol2* background, TSO1 helps ensure the division of the GMC prior to differentiation.

We quantified SGC phenotypes in two independent *sol1 sol2; amiRNA-tso1* lines and confirmed that SGCs were unique to this triple depletion genotype (Fig. 6E). In doing so, we also noticed that *sol1 sol2; amiRNA-tso1* had fewer stomatal pairs and that the stomata and pavement cells were visibly larger than WT or *sol1 sol2* (Fig. 6D). These phenotypes were opposite that of *sol1 sol2* alone; therefore we asked whether depletion of *TSO1* could “rescue” the stomatal pairing and small cell phenotypes associated with loss of *SOL1* and *SOL2*. When quantified, the *sol1 sol2; amiRNA-tso1* lines had fewer cells per field of view than *sol1 sol2* plants (Fig. S5A). We normalized the number of stomatal pairs to the number of pavement cells per field of view and found the number of pairs was still reduced in amiRNA-tso1 *sol1 sol2* lines compared to *sol1 sol2* mutants (Fig. 6F). The rescue of the *sol1 sol2* pairing phenotype, as well as the larger pavement cells and guard cells suggested a repression of cell division in the epidermis.

The phenotypic effects on stomatal lineage cells suggested that *TSO1* acts in opposition to *SOL1* and *SOL2*. To test this idea further, we overexpressed SOL2, reasoning that more SOL2 would produce same SGC phenotype as loss of *TSO1*. We placed SOL2-CFP under the control of a strong, estradiol inducible promoter and induced 3 dpg seedlings bearing the transgene with estradiol for 8 hours, monitored expression of CFP to confirm overexpression of SOL2 (Fig. S5B), then returned seedlings to plates to grow for an additional 5 days. The SOL2-overexpressing seedlings produced SGCs (Fig. 6H, blue arrowhead), whereas the equivalent estradiol treatment on a control line did not (Fig. 6G). The majority of SOL2-CFP expressing seedlings exhibited SGCs on both the adaxial and abaxial surfaces (Fig. S5C). We concluded that at the GMC stage of stomatal lineage development, three closely related CHC proteins could have opposite effects on cell cycle progression, with TSO1 acting is a positive regulator and SOL2 (and SOL1) as negative regulators.

## Discussion

As a key regulator of the stomatal lineage, SPCH activates and represses thousands of genes to start the proliferative meristemoid phase of the lineage. Logically, SPCH must also set in place a program that will allow cells to exit this proliferative stage. SPCH directly activates many of its own negative regulators, including BASL, EPF2 and TMM, suggesting the existence of feedback loops that modulate SPCH levels (Horst et al., 2015; Lau et al., 2014). Here we have shown that *SOL1* and *SOL2* are stomatal lineage expressed SPCH transcriptional targets and that they encode proteins with a distinctive cycling expression pattern (Fig. 7A). Normally when cells stop expressing SPCH they either begin expressing MUTE and transition to GMC fate, or they become SLGCs and differentiate into pavement cells. Our data suggest that SOL1 and SOL2 aid SPCH-expressing meristemoids in their timely transitions to either of these later fates. For example, time-lapse imaging of cell fate reporters in *sol1 sol2* mutants revealed that MUTE-expressing cells could still have the division behaviors associated with SPCH-expressing cells, whereas other SPCH-expressing cells fail to differentiate morphologically into pavement cells even after they downregulate SPCH. How might SOL1 and SOL2 aid in transitions? One possibility is that, as DNA-binding domain containing proteins, they regulate expression of *SPCH*. In support of this idea, a genome-wide analysis of Arabidopsis transcription factor binding found SOL1 and SOL2 associated with sequences immediately upstream of *SPCH* (O’Malley et al., 2016). Alternatively, SOLs might repress meristemoid identity genes downstream of SPCH when that phase ends.

**Figure 7:**
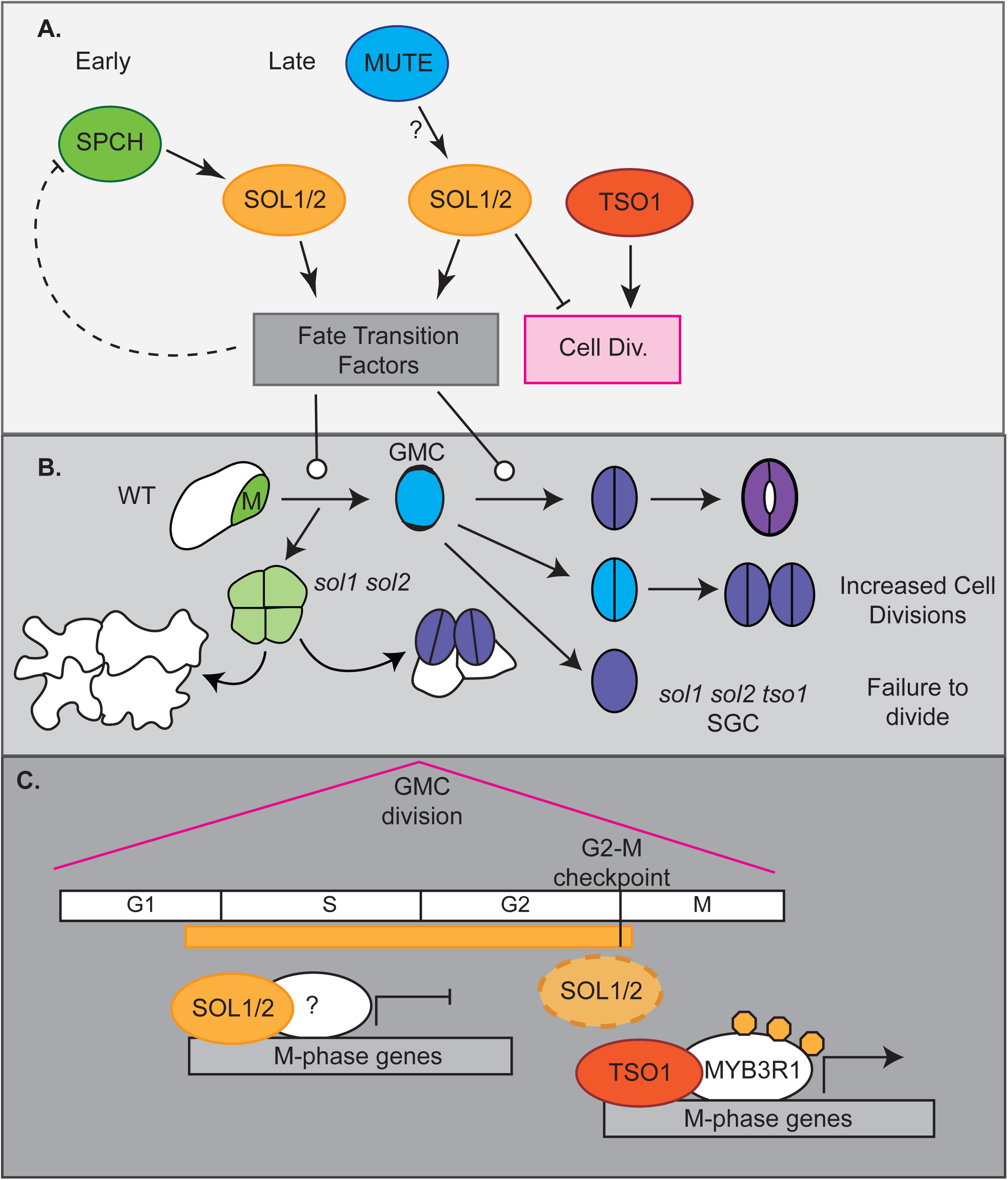
A model of SOL function in stomatal fate transitions and cell divisions. **(A)** In meristemoids, SPCH binds to and induces *SOL1* and *SOL2,* and their protein products regulate the M→GMC transition and may downregulate *SPCH* in a negative feedback loop. In GMCs, MUTE induces *SOL1* and *SOL2* to regulate the GMC→GC transition and limit cell divisions. At this stage, SOL1 and SOL2 oppose TSO1. **(B)** In *sol1 sol2* mutants, meristemoids fail to progress to SLGC or GMC identity in a timely manner, although they may eventually become stomata (sometimes forming pairs) or pavement cells. Therefore, stomatal pairs arise from two different defects in fate transition – 1 early and 1 late. In the absence of *tso1* GMCs fail to divide forming single guard cells (SGC). **(C)** SOL1 and SOL2 repress divisions, possibly by repressing M-phase genes in S and G2. In M-phase, SOL1 and SOL2 disappear and TSO1 is able to upregulate M-phase genes through its binding partner, MYB3R1.

Our analysis of the expression pattern and mutant phenotype also revealed roles of SOL1 and SOL2 at post-SPCH stages of stomatal development. Interestingly, a recent study found that both genes are upregulated in response to MUTE induction (log2 fold changes of 1.60 and 0.83 respectively) (Han et al., 2018). Whether these genes are direct MUTE targets is not known, but the appearance of SOL1 in GMCs shortly following MUTE expression (Fig. 2J-K) is consistent with it being a MUTE target. The broader expression pattern of *SOL2* suggests it is likely dependent on other inputs, consistent with the weaker induction of *SOL2* relative to *SOL1* in both SPCH and MUTE induction experiments (Han et al., 2018; Lau et al., 2014). The inappropriate expression of MUTE in small cells may suggest that SOL1 and SOL2 downregulate MUTE in a negative feedback loop; however, neither SOL1 nor SOL2 was found to bind upstream of *MUTE* in large-scale assays of transcription factors (O’Malley et al., 2016). Alternatively, as downstream targets of MUTE, SOL1 and SOL2 could be coordinating divisions with fate transitions (Fig. 7B). In this model, MUTE is expressed in the small cells at the correct time, but in the absence of SOL1 and SOL2, these cells fail to transition to GMCs and continue to undergo meristemoid-like divisions.

*SOL1, SOL2* and their paralogue *TSO1,* which is not a direct target of SPCH, but is nonetheless expressed in the epidermis, are then involved in the next fate transition from GMC to guard cell. In wildtype, this transition is tied to the symmetric division of the GMC into two guard cells. In *sol1 sol2* mutants, ectopic GMC-like divisions of young guard cells can result in stomatal pairs. Overexpression of SOL2 or knockdown of *TSO1* in the *sol1 sol2* background leads to the opposite phenotype in which GMCs fail to divide, suggesting oppositional roles of SOL1/2 and TSO1 at the GMC division (diagrammed in Fig. 7B). Cell fate is intrinsically tied to cell division; therefore, it is not always possible to cleanly separate the two. For example, loss of FAMA expression leads to immature guard cells that recapitulate GMC divisions (Ohashi-Ito and Bergmann, 2006). If SOL1 and SOL2 promote differentiation, then in their absence young guard cells retain GMC fate long enough to divide a second time. In the absence of *tso1 sol1* and *sol2*, GMCs differentiate and lose the ability to divide too quickly, resulting in SGCs. However, these proteins might also directly alter the cell cycle (Fig. 7C).

We cannot ignore the distinct cell cycle expression pattern of the SOLs, especially in light of the cell cycle regulatory role that animal CHC domain containing proteins play. In animals, which typically encode a single somatic CHC domain-containing protein, the CHC protein is found in two types of DREAM complexes: the quiescent DREAM complex whose role is to repress gene expression in G_0_ and the MYB-containing “permissive” DREAM complex found in actively proliferating cells (Beall et al., 2004; Beall et al., 2007). DREAM also regulates gene expression and epigenetic marks outside of the cell cycle, for example it regulates the expression of olfactory receptors in fly neurons via histone methylation (Sim et al., 2012).

CHC proteins in plants could potentially be activators or repressors of the cell cycle, and function in both MYB and DREAM dependent and independent ways. We suggest that SOL1 and SOL2 would have a cell cycle repressive role based on the observations that (1) *sol1 sol2* mutants have leaves with more cells (2) *sol1 sol2* mutants display inappropriate divisions of MUTE-expressing cells and (3) SOL1 and SOL2 proteins disappear prior to cell division. In contrast, *TSO1* mutants, originally identified by reproductive development defects, have cytokinesis defects, shorter root apical meristems and are sterile (Andersen et al., 2007; Hauser et al., 2000; Liu et al., 1997; Sijacic et al., 2011), which suggest that TSO1 promotes divisions, though fasciation of the floral meristem in *tso1* indicates *TSO1* may restrain divisions in some contexts. *Arabidopsis* encodes many proteins with MYB domains, including FOUR LIPS (FLP) and its closest paralogue MYB88, which have been connected to GMC divisions (Lai et al., 2005). The *Arabidopsis* MYBs most likely to be involved in a DREAM complex, however, are five three-repeat-MYB proteins (MYB3R1-5) more structurally similar to the animal MYBs than FLP and MYB88 (Stracke et al., 2001). Recent work has linked TSO1 to activity of MYB3R1 a MYB with both cell cycle activating and repressive roles (Araki et al., 2004; Ito et al., 2001; Kobayashi et al., 2015). Mutations in *MYB3R1* suppress the *tso1-1* phenotype and TSO1 physically interacts with MYB3R1 (Wang et al. 2018). Moreover, SGC phenotypes have been reported in *myb3r1 myb3r4* double mutants (Haga et al., 2007); although it is not known whether the SGC phenotypes in these mutant backgrounds all arise from defects in the same stage of the cell cycle.

These discoveries, along with evidence of other physical interactions between DREAM complex homologues (for example, SOL1 appeared as a partner of repressive MYB3R3 in a proteomics-based analysis (Kobayashi et al. 2015)), has led to the hypothesis that CHC proteins function in a plant version of the DREAM complex (reviewed in Magyar and Ito, 2016). How might we imagine a DREAM complex acts in the stomatal lineage? Perhaps our most unexpected finding was that SOL1 and SOL2 expression patterns overlap their homologue TSO1 in the epidermis, but phenotypes associated with their loss or overexpression are opposite. This is a novel situation for DREAM complexes as there are only single CHC and MYB proteins available for the animal somatic complexes. The function of MYB3R1 as both activator and repressor of cell cycle progression adds another layer of complexity. Phosphorylation state may contribute to its dual function (Araki et al., 2004; Chen et al., 2017; Wang et al., 2018), however, binding partners could also play a role.

An earlier model postulated that TSO1 interacts with MYB3R1 to drive M-phase gene activation (Fig. 7C)(Kobayashi et al., 2015; Wang et al., 2018). Given that SOL1 can interact with the repressive MYB3R3, we can imagine several additions to that core model. SOL1 and SOL2 might interact with repressive MYBs to limit the expression of M-phase genes, but their disappearance from dividing cells 1-2 hours before the appearance of the new cell plate, could be part of a G2-M switch mechanism, in which proteolytic degradation of SOL1/2 leads to incorporation of TSO1 and the activator MYBs into a plant DREAM complex. An alternative hypothesis is that SOL1, SOL2 and TSO1 can all interact with both types of MYB3Rs. In this model, MYB3R1 switches from a repressor to an activator when SOL1 and SOL2 are degraded at G2-M and instead it binds to TSO1. When SOL2 is overexpressed, it sequesters the MYB3R1 protein in the repressor complex, recapitulating the *sol1 sol2 amiR-tso1* phenotype and the *myb3r1 myb3r4* phenotype. Similarly, in *sol1 sol2*, only the MYB3R1-TSO1 activating complex is present leading to inappropriate divisions. Finding the precise molecular mechanism for the diverse CHC family roles in cell behaviors will be an intriguing but challenging future goal, as it will require quantitative assays of differential incorporation of CHCs into functional complexes, coupled to measurements of gene expression in response to different complexes in the relevant cell types.

Key regulators of three separate stomatal cell states have been known for many years; here we add an important feature to the developmental trajectory: CHC-domain proteins to enforce transitions between these fates and to regulate their associated cell cycle behaviors. New technologies enabling measures of transcriptomes and chromatin accessibility in individual cells have reinvigorated the idea of “transitional states”, and while there are computational methods to identify where and when these states occur (Farrell et al., 2018; Xiao et al., 2018) how they are resolved will require experimental analysis of regulators like the SOLs. We focused on the stomatal lineage, and found multiple fate transitions are regulated by the same factors, leading to the interesting possibility that CHC proteins and the DREAM complex will be used repeatedly for cell fate transitions in other tissue, organs and stages of plant development.

## MATERIALS AND METHODS

### Plant material and growth conditions

Arabidopsis thaliana Columbia (Col-0) was used as wild type in all experiments. Seedlings were grown on half-strength Murashige and Skoog (MS) medium (Caisson Labs) at 22°C in an ARR66 Percival Chamber under 16 h-light/8 h-dark cycles and were examined at the indicated times. The following previously described mutants and reporter lines were used in this study: SPCHpro:SPCH-CFP and MUTEpro:MUTE-YFP (Davies and Bergmann, 2014); FAMAproYFPnls (Ohashi-Ito and Bergmann, 2006); HTR2pro:CDT1a(C3)-RFP (Yin et al., 2014); TSO1pro:TSO1-GFP (Wang et al., 2018); *tso1-5* (Salk_102956)(Andersen et al., 2007), *hdg2-2*(SALK_127828C) and *hdg2-4*(SALK_120064)(Peterson et al., 2013). The following lines were obtained from the ABRC stock center: *sol1-3*(SAIL_742_H03), *sol1-4* (WiscDsLoxHs033_03E), *sol2-2* (SALK_021952), *sol2-3* (SALK_031643). The HDG2proHDG2-GFP construct (Peterson et al., 2013) was a kind gift from Prof. Keiko Torii (University of Washington)

### Vector construction and plant transformation

Constructs were generated using the Gateway system (Invitrogen). Appropriate genome sequences (PCR amplified from Col-0 or from entry clones) were cloned into Gateway-compatible entry vectors, typically pENTR/D-TOPO (Life Technologies), while promoter sequences were cloned into pENTR-5’TOPO (Life Technologies) to facilitate subsequent cloning into plant binary vectors pHGY (Kubo et al., 2005) or R4pGWB destination vector system (Nakagawa et al., 2008).

Transcriptional reporters for *SOL1* and *SOL2* were generated by cloning a 5’ regulatory region spanning 2500bp or to the 3’ end of the upstream gene or (whichever was shorter) to the ATG translational start site into pENTR5’ and recombining with pENTR YFP into R4pGWB540 (Nakagawa et al., 2008). For the SOL1 and SOL2 translational fusions, the genomic fragments corresponding to *SOL1* and *SOL2* (excluding stop codon) were amplified by PCR then cloned in pENTR D/TOPO (Life Technologies) LR Clonase II was then used to recombine the resulting pENTR clone and pENTR 5’ promoters (SOL1p, SOL2p) into R4pGWB540. For the estradiol inducible lines, the UBQ10 promoter was amplified by PCR and subcloned into pJET, then digested out using AscI XhoI double digest and ligated into p1R4:ML-XVE (Siligato et al., 2016). P1R4:UBQ10-XVE was recombined with SOL2 pENTR and R4pGWB443 (Nakagawa et al., 2008). The TSO1 amiRNA was generated as described previously (Sijacic et al., 2011).

Transgenic plants were generated by Agrobacterium-mediated transformation (Clough, 2005), and transgenic seedlings were selected by growth on half-strength MS plates supplemented with 50 mg/l Hygromycin (pHGY-, p35HGY-, pGWB1-, pGWB540-based constructs), 100 mg/l Kanamycin (pGWB440 based constructs) or 12 mg/l Basta (pGWB640-based constructs). Primer sequences used for entry clones are provided in Table S1.

### Estradiol induction

3 dpg seedlings grown on agar-solidified half strength MS media were flooded with 10 uM estradiol (Fluka Chemicals) or a vehicle control. At 8 hrs post induction, liquid was removed, and plates were allowed to dry, before being returned to incubator for 5 more days. Tissue was collected at 8 dpg and cleared in 7:1 Ethanol:Acetic acid.

### Confocal and differential interference contrast microscopy

For confocal microscopy, images were taken with a Leica SP5 microscope and processed in ImageJ. Cell outlines were visualized by 0.1 mg/ml propidium iodide in water (Molecular Probes). Seedlings were incubated for 10 min in the staining solution and then rinsed once in H2O. For differential interference contrast (DIC) microscopy, samples were cleared in 7:1 ethanol:acetic acid, treated for 30 min with 1N potassium hydroxide, rinsed in water and mounted in Hoyer’s medium. DIC images were obtained on a Leica DM2500.

### Statistical Analysis

Image J was used to count clustering events within a defined field of view. Statistical analysis was completed in Graphpad Prism. For clustering and cell counts, data were generally not normally distributed (based on D’Agostino-Pearson test) so analysis was completed with default settings for nonparametric tests. The Mann-Whitney test was used, where indicated, to compare two sets of data; to compare multiple groups against one another, the Kruskal-Wallis test, followed by Dunn’s multiple comparison test was used where indicated in figure legends.

### RT-qPCR analysis

RNA was extracted from 9 dpg whole seedlings (*sol1-3, sol1-4, sol2-2, sol2-3* and *sol1-4 sol2-2* double mutants, and WT controls) using the RNeasy Plant Mini Kit (Qiagen) with on-column DNAse digestion. cDNA was synthesized with iSCRIPT cDNA Synthesis Kit (BioRAD), followed by amplification with the SsoAdvanced™ SYBR® Green Supermix (Bio-Rad) using gene specific primers on a CFX96 Real-Time PCR detection system (Bio-Rad). Reaction conditions: Data were normalized to *ACTIN2* gene controls using the ΔΔ^CT^ method. Three biological replicates were assayed per genotype. Primers are listed in Table S1.

### Time-lapse imaging

After growth on half strength MS media, seedlings were transferred to a sterilized perfusion chamber at indicated days post germination for imaging on a Leica SP5 Confocal microscope following protocols described previously (Davies and Bergmann, 2014). The chamber was perfused with 1/4 strength.75% (w/v) sucrose (or glucose) liquid MS growth media (pH 5.8) at a rate of 2mL/hr. Z-stacks through the epidermis were captured with Leica software every 30 or every 60 minutes over 12-60 hour periods and then processed with Fiji/ImageJ (NIH). Areal growth calculated by determining the 2D area immediately after one division (Area1) and immediately prior to the next division of the same cell (Area2) using ImageJ.

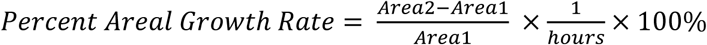

## Acknowledgements

We thank lab members Nathan Cho for constructing the SOL2-YFP, CDT1a-RFP line, Yan Gong and Dr. Heather Cartwright (Carnegie) for imaging advice, and Dr. Annika Weimer and Dr. Camila Lopez-Anido for detailed feedback on the manuscript.

## Author contributions

Conceptualization: A.R.S, K.A.D, D.C.B.; Methodology: ARS, KAD Formal analysis: ARS, KAD; Investigation: ARS, KAD; Resources: W.W. Z.L. Writing - original draft: A.R.S, K.A.D, D.C.B; Writing - review & editing: A.R.S, W.W. Z.L. K.A.D, D.C.B; Visualization: A.R.S, K.A.D; Supervision: D.C.B.; Project administration: D.C.B.; Funding acquisition: D.C.B.

## SUPPLEMENTARY INFORMATION

Figure S1: Additional analysis of SOL1 and SOL2 expression patterns emphasizing cell cycle expression

Figure S2: Supporting information about alleles used for phenotypic analysis

Figure S3: Evidence that cell cycle times are increased, and post-division cell growth reduced in the stomatal lineage of *sol1 sol2* plants

Figure S4: Additional marker in *sol1 sol2* double mutants and marker expression in wildtype seedlings.

Figure S5: Quantification of effects of *tso-1* amiRNA and SOL2-CFP overexpression on cell size and division phenotypes

Table S1: Primers used in this study

**Figure S1:**
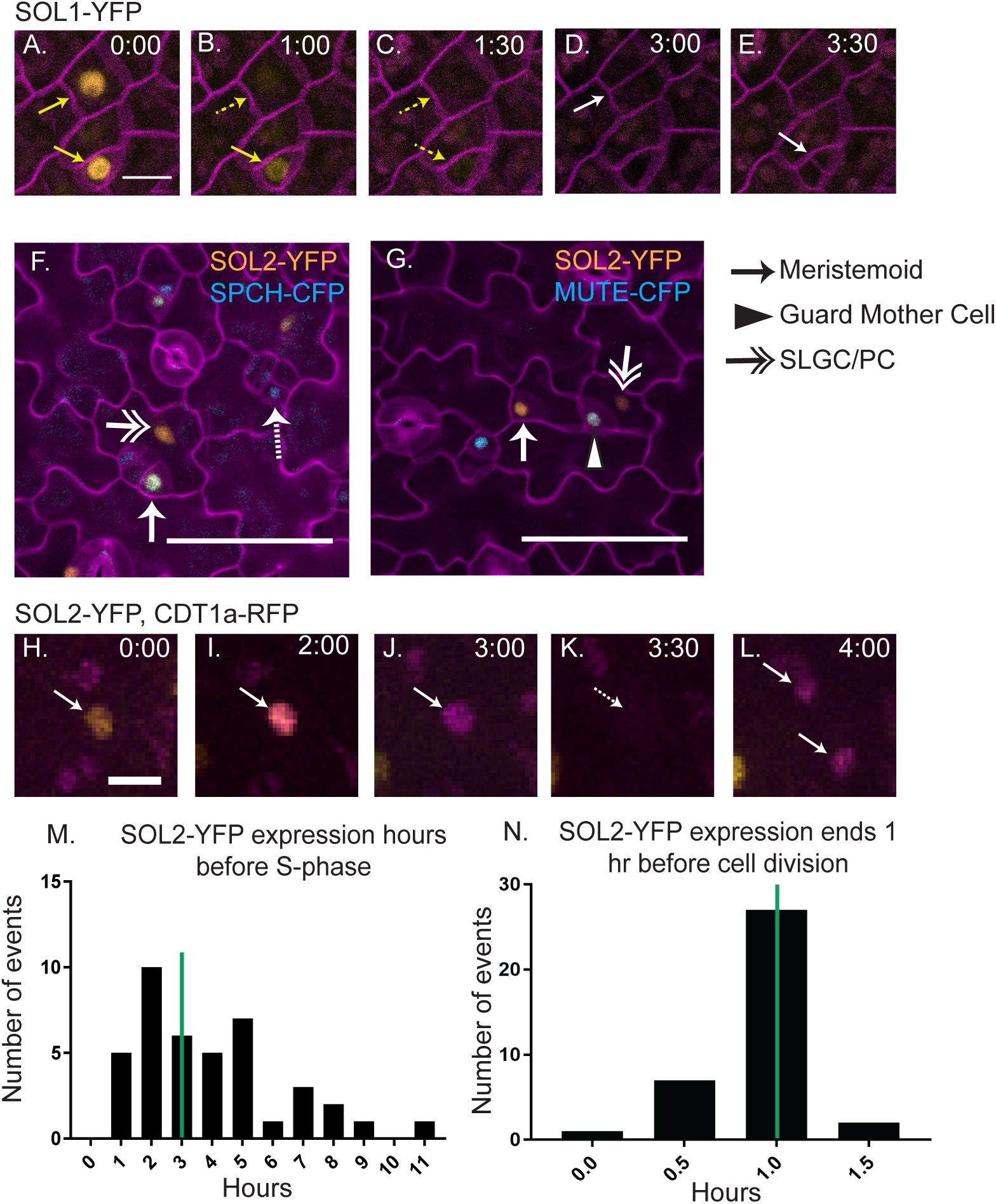
Additional analysis of SOL1-YFP and SOL2-YFP, emphasizing connections to cell cycle. (A-E) Time-lapse confocal imaging of SOL1pro:SOL1-YFP in wildtype background; plasma membrane visualized with ML1pro:RCI2A-mCherry, image captured every 30 min. SOL1 is expressed in two cells (A, yellow arrows). It turns off in the upper cell (B, dotted yellow arrow) then the lower cell (C, dotted yellow arrow). Each cell divides 2 hrs after SOL1-YFP expression is last seen (D, upper cell, white arrow) (E, lower cell, white arrow). (F-G) SOL2pro:SOL2-YFP is co-expressed with SPCHpro:SPCH-CFP in some (white arrow), but not all meristemoids (white dotted arrow) and with MUTEpro:MUTE-CFP in GMCs (arrowhead). SOL2 is also expressed in pavement cells and SLGCs (double arrows) that don’t express SPCH or MUTE. (H-L) Representative images from time-lapse of SOL2pro:SOL2-YFP, HTR2pro:CDT1a(C3)-RFP. SOL2-YFP is visible first (H), then co-expressed with CDT1a-RFP (I). CDT1a-RFP is not visible for one frame (K) presumably during nuclear envelope breakdown, however, it persists into both daughter cells (L). (M) Quantification of length of time that YFP is detected before RFP is detected, green line indicates median at 3 hours, n=41. (N) Quantification of length of time after YFP cannot be seen before cell division, n=37, green line indicates median at 1 hour.

**Figure S2:**
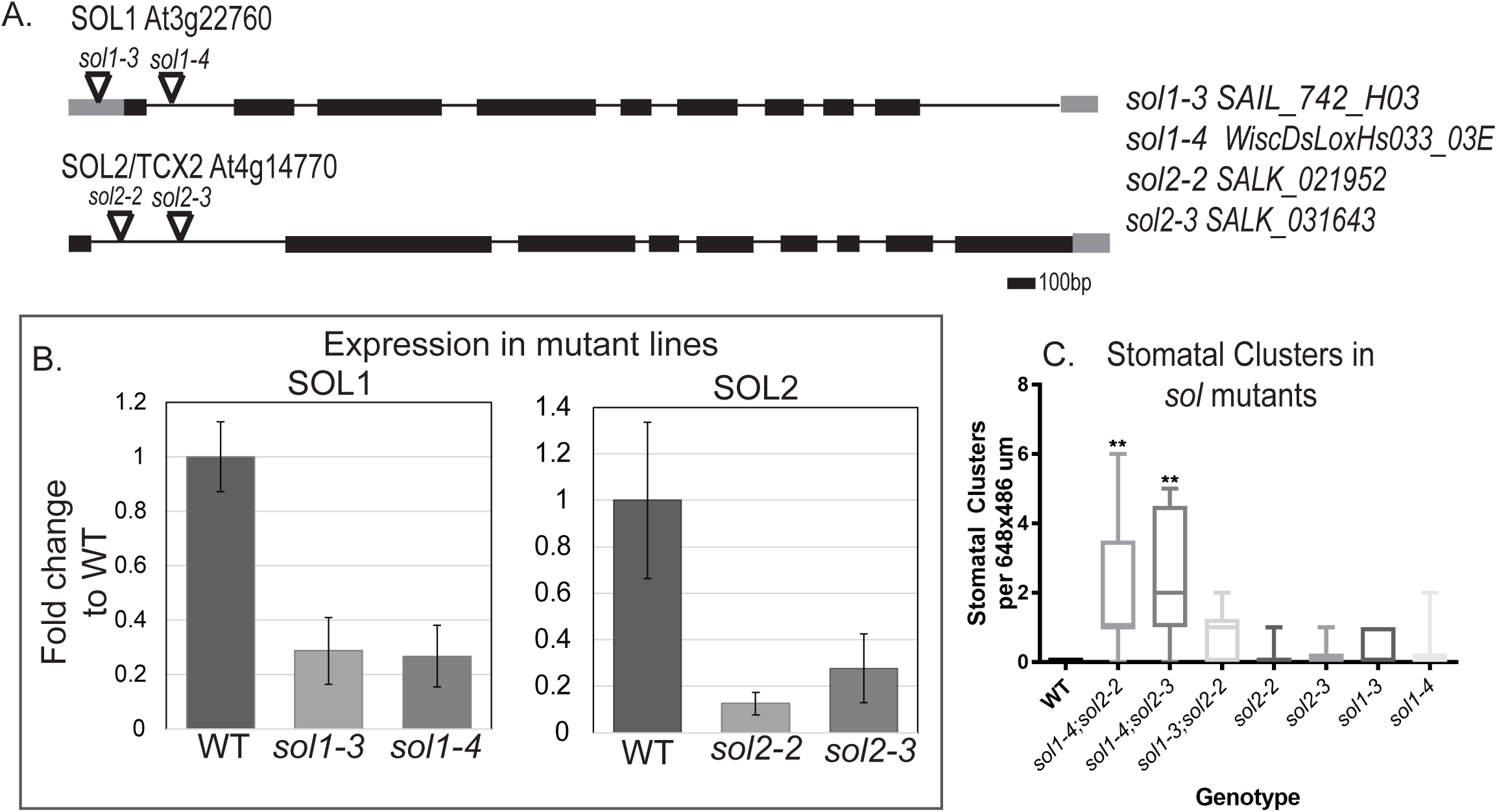
Supporting information about alleles used for phenotypic analysis. (A) Diagram of SOL1 and SOL2 genomic loci with position of T-DNA alleles indicated by triangles. (B) qRT-PCR analysis of expression levels of SOL1 and SOL2 transcripts in mutant seedlings at 9 dpg, levels are normalized to ACT2 as a reference gene, 3 biological replicates per genotype, error bars indicate standard deviation. (C) Quantification of stomatal clusters phenotypes in SOL single and double mutants, n = 9-10, significant difference compared to WT ** p<0.01, Dunn’s multiple comparison test.

**Figure S3:**
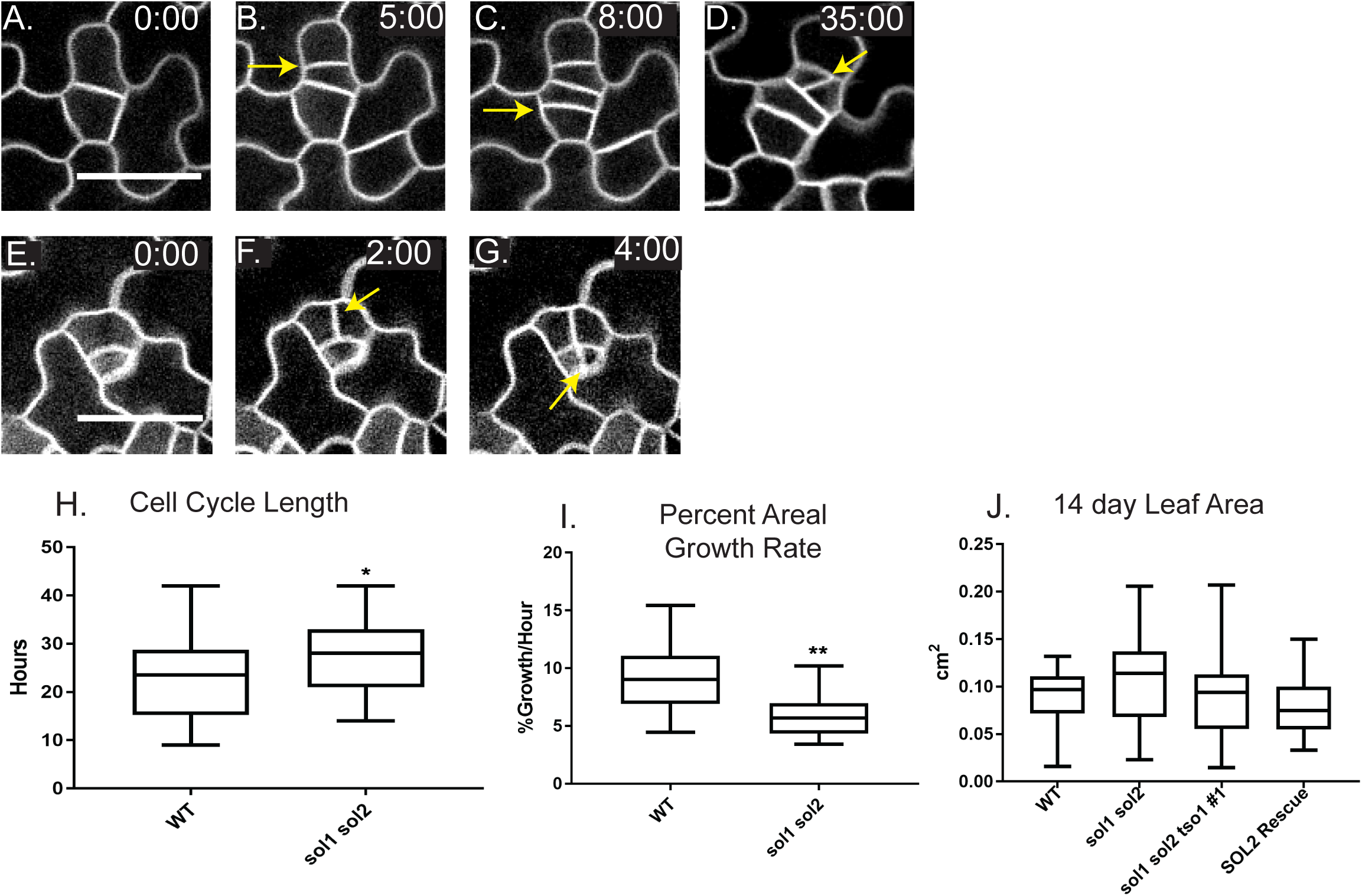
Evidence that cell cycle times are increased, and post-division cell growth reduced in the stomatal lineage of *sol1 sol2* plants. (A-G) Confocal time-lapse images of cells dividing in *sol1 sol2* as an example of data quantified in H-J, divisions indicated with yellow arrows. Scale bar 30 μm. (H) Cell cycle length is increased in *sol1 sol2* mutants (WT n=24 cells scored, sol1 sol2 n=22). (I) Percent growth per hour in small cells is reduced in *sol1 sol2* mutants (WT n=14 cells scored, sol1 sol2 n=13). (J) Overall true leaf area at 14 dpg is not significantly different between WT and *sol1 sol2* mutants. Significance indicated: * p<0.05, ** p<0.01, *** p<0.001, Mann Whitney test.

**Figure S4:**
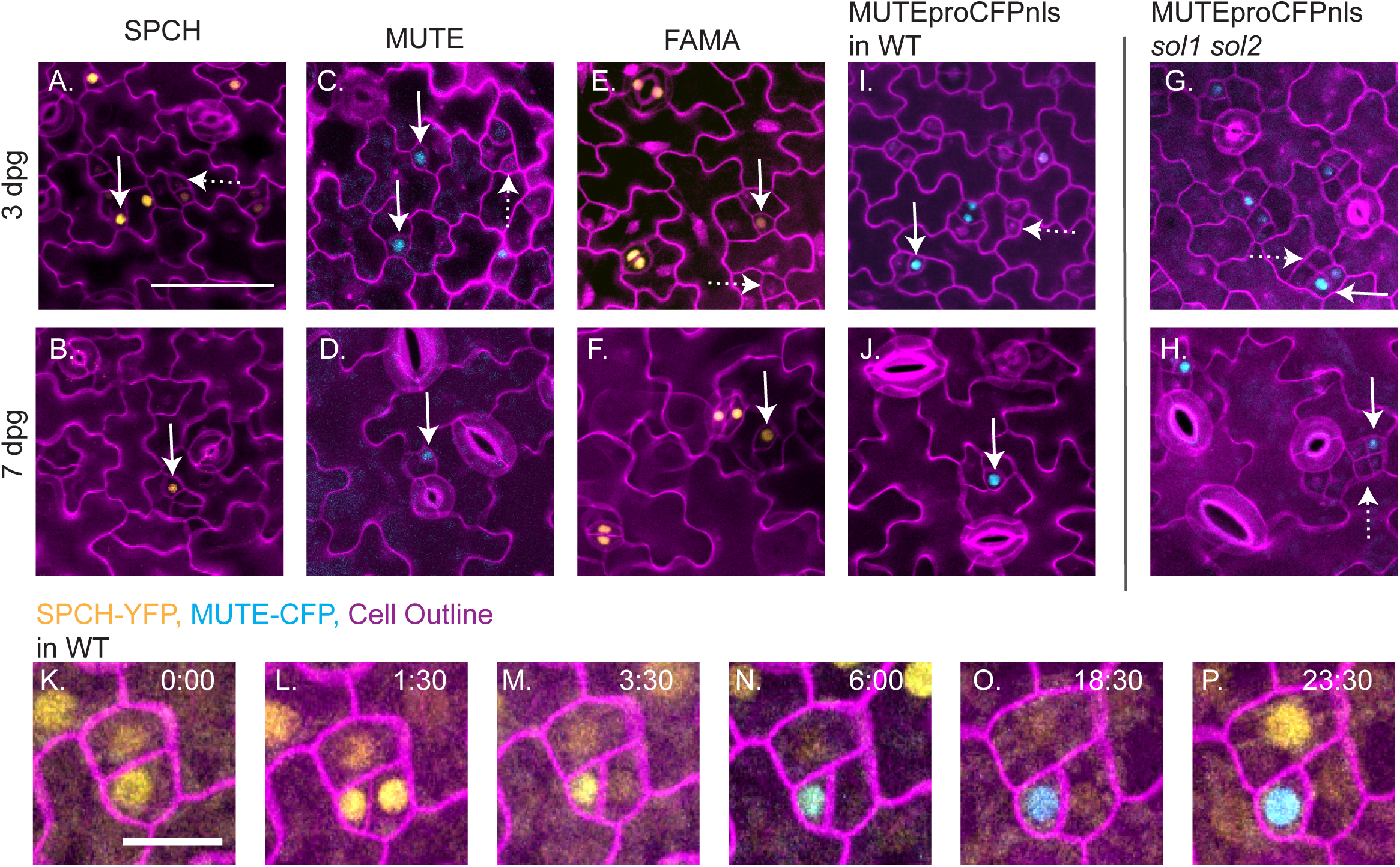
Additional marker in sol1 sol2 double mutants and marker expression in wildtype seedlings. (A-B) SPCHpro:SPCH-YFP in wildtype seedlings. (C-D) MUTEpro:MUTE-CFP in wildtype seedlings. (E-F) FAMApro:YFPnls in wildtype seedlings. (I-J) MUTEpro:CFPnls in wildtype seedlings. (G-H) MUTEpro:CFPnls in sol1 sol2 seedlings. All images at same scale, scale bar in A, 50 μm. (K-P) Selections from time-lapse of ML1pro:RCI2A-mCherry, SPCHpro:SPCH-YFP and MUTEpro:MUTE-CFP markers, all images same scale, scale marker in (K) 20 μm. SPCH expressing cell divides in (L), begins to express MUTE in (N).

**Figure S5:**
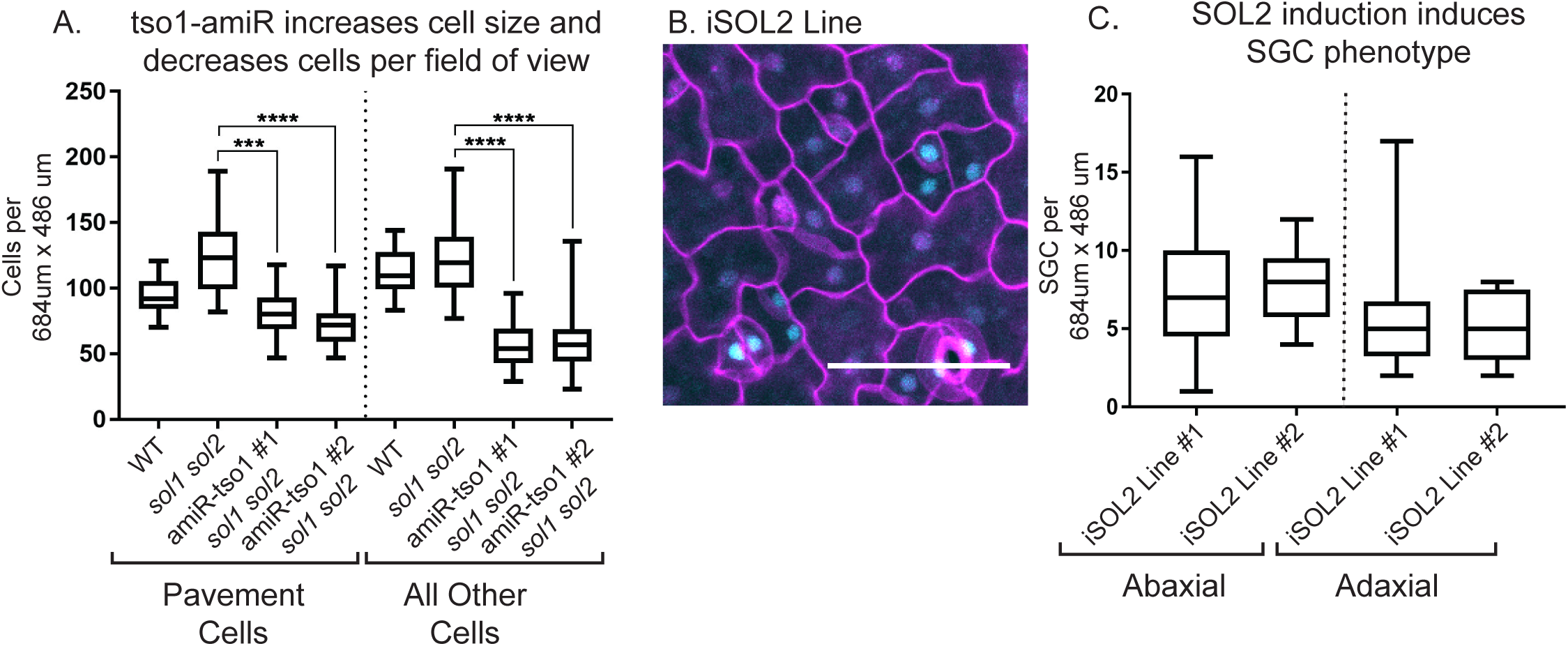
Quantification of effects of *tso-1* amiRNA and SOL2-CFP overexpression on cell size and division phenotypes. (A) Quantification of the changes in cell size and numbers in *tso1-amiRNA sol1 sol2* shows a decreased number of pavement cells and other cells (non-pavement cells, including guard cells) relative to *sol1 sol2*. (B) Expression of SOL2-CFP in 4dpg seedling throughout epidermis 24 hours after beta-estradiol induction. (C) Incidence of SGCs per field of view in two independent lines of induced seedlings. Seedlings induced at 3 dpg, screened for expression, then collected for analysis at 8 dpg, n = 9-13. Significance indicated: *** p<0.001, **** p<0.0001, Dunn’s multiple comparison test.

**Table S1:**
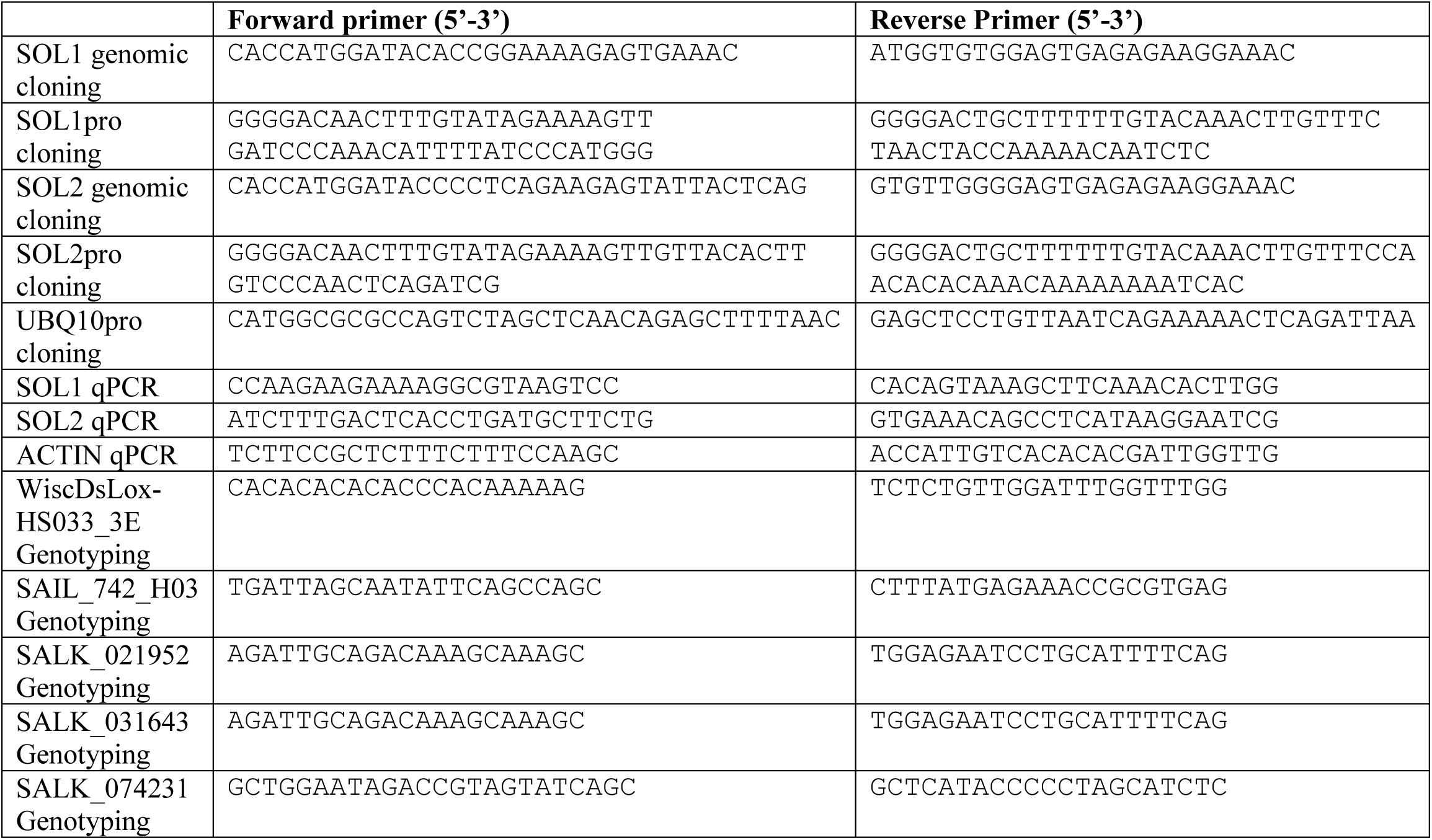
Primers used in this study

